# 3D Quantification of Viral Transduction Efficiency in Living Human Retinal Organoids

**DOI:** 10.1101/2024.03.06.583795

**Authors:** Teresa S. Rogler, Katja A. Salbaum, Achim T. Brinkop, Selina M. Sonntag, Rebecca James, Elijah R. Shelton, Alina Thielen, Roland Rose, Sabrina Babutzka, Thomas Klopstock, Stylianos Michalakis, Friedhelm Serwane

**Author notes:** **Declaration of Interests** S.M. is cofounder of the gene therapy company, ViGeneron GmbH, who owns the rights on the related patent application WO/2019/076856 covering the AAV2.NN and AAV9.NN. All other authors declare no competing interests.

## Abstract

The development of therapeutics builds on testing their efficiency in vitro. To optimize gene therapies, for example, fluorescent reporters expressed by treated cells are typically utilized as readouts. Traditionally, their global fluorescence signal has been used as an estimate of transduction efficiency. However, analysis in individual cells within a living 3D tissue remains a challenge. Readout on a single-cell level can be realized via fluorescence-based flow cytometry at the cost of tissue dissociation and loss of spatial information. Complementary, spatial information is accessible via immunofluorescence of fixed samples. Both approaches impede time-dependent studies on the delivery of the vector to the cells.

Here, quantitative 3D characterization of viral transduction efficiencies in living retinal organoids is introduced. The approach combines quantified gene delivery efficiency in space and time, leveraging human retinal organoids, engineered adeno-associated virus (AAV) vectors, confocal live imaging, and deep learning-based image segmentation. The integration of these tools in an organoid imaging and analysis pipeline allows quantitative testing of future treatments and other gene delivery methods. It has the potential to guide the development of therapies in biomedical applications.

## Introduction

AAV (adeno-associated virus) vectors are a common delivery vehicle for gene therapy applications. AAV-based gene therapies are effective in various tissue types, such as the retina, and are being tested in several clinical studies.^[1,2]^ In 2017, the first gene therapy for retinal degenerative diseases, Luxturna, was approved for the markets in the US and European Union^[3]^ To assess key properties like viral serotype transduction efficiency, pervasion depth, and systemic effects, a reliable testing platform is essential, ideally in human tissue.

Human retinal organoids (hROs), which are stem cell-derived in vitro models for human retinal tissue, have the potential to transform preclinical drug testing.^[4–6]^ State-of-the-art protocols enable the high throughput culturing of hROs that possess tissue-specific cell types, morphological characteristics of the neural retina (NR), and the retinal pigment epithelium (RPE). hROs display functional properties of the retina light sensitivity and synaptic connectivity ^[7–11]^ and thus enable insights into the physiological mechanisms behind retinal development ^[12,13]^ and diseases.^[14]^ Recent studies used hROs to test and optimize mRNA delivery via liposomes ^[15]^ and gene delivery via AAV vectors.^[1,2,16–18]^ Hence, they are now emerging as a complementary in vitro system for gene therapy testing.^[19,20]^

Transduction efficiency of novel AAV vectors is typically assessed with vector genomes coding for fluorescent proteins. The fluorescence expression is then used as an indicator of the vector’s ability to transduce cells. Traditionally, estimates of transduction efficiencies were either obtained by recording fluorescence levels without further quantification,^[2]^ normalizing the overall fluorescence level of a whole organoid to that of a constitutive nuclear stain like DAPI,^[17]^ or comparing the overall fluorescence level obtained with different vectors without normalization.^[16]^ These approaches face the fundamental challenge that a single transduced cell with strong fluorescence reporter expression cannot be distinguished from multiple transduced cells with low fluorescence levels (Figure 1). This ambiguity makes the assessment of viral transduction efficiency difficult. To quantify transduction on a single-cell level, fluorescence-based flow cytometry can be performed.^[18]^ However, due to the requirement of organoid dissociation, spatial information is lost and signals from non-retinal tissue, as found in retinal organoids,^[7]^ cannot be excluded from the single-cell pool. To access spatial transduction information, researchers typically use fixed samples ^[22]^ which limits readout of critical information, such as viral propagation, that can be gained through dynamic longitudinal studies on the same sample.

**Figure 1.**
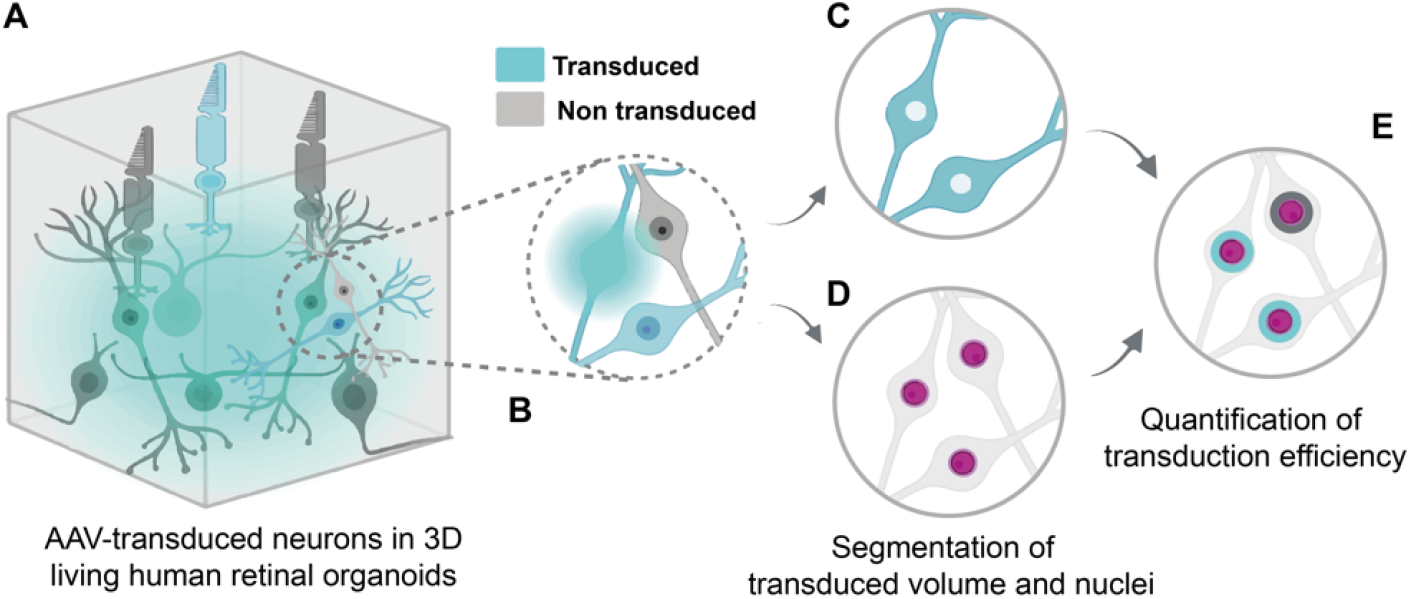
Quantification of viral vector transduction efficiencies in living human retinal organoids. (A) Cells are densely packed and (B) show varying expression levels of the transgene if transduced. A quantitative measure of the transduction efficiency is obtained by segmentation of the transduced volume (C) via machine learning and of the nuclei (D) via deep learning techniques.^[21]^ Transduced cells are classified by evaluating the transduction segmentation within a thin virtual shell around each nucleus (E).

Here, we introduce 3D quantification of AAV transduction efficiency in living retinal organoids with single-cell resolution (Figure 1). To sample the entire retinal cross-section of living hROs, we imaged their retinal part using confocal microscopy (Figure 1A). The segmentation of individual neurons in a densely packed tissue has previously required specialized imaging techniques like super-resolution microscopy, which poses severe challenges to the imaging hardware.^[23]^ To lower the hurdle and still obtain single cell resolution, we focused on the segmentation of the overall transduced volume instead of separation of individual cells. To classify this volume independently of the fluorescence signal intensity, we used a machine learning algorithm which we trained with a minimal set of data consisting of transduced cells with varying brightness (Figure 1C, see Methods). To segment cell nuclei, we used the deep learningbased segmentation software Cellpose (Figure 1D) due to its performance in dense tissues.^[21]^ To evaluate whether a cell was transduced, we quantified the overlap of a thin shell covering each nucleus with the transduced volume (Figure 1E). We used our pipeline to test three AAVs with different capsids (AAV2.7m8, AAV9.NN, AAV2.NN), and tracked their transduction efficiency spatially over a period of 40 days with four individual time points. The ability to quantify transduction efficiencies in living organoids enables preclinical evaluation and optimization of viral vectors based on their performance in human retinal tissue. Moreover, it opens the door to spatially resolved longitudinal studies for any delivery experiment such as electroporation or cell lines including cytoplasmic tissue-specific fluorescent reporters.

## Results

### Culture of human retinal organoids

We grew human retinal organoids according to the protocol by Cowan *et al.* with minor adjustments (see Experimental Section)^[7]^ As a starting point, we used induced pluripotent stem cells (iPSCs) to generate embryoid bodies, differentiated them towards eye field fate in an attached state (2.5D) and let them mature into retinal organoids within a suspended environment (Figure 2A-D). The iPSCs and organoids were tested for pluripotency and retina-specific cell types by immunostaining to confirm the cell fate (Supplementary Figure S1-2; Supplementary Table S1). The mature organoids were then used for the experimental setup to determine the transduction efficiency within confocal 3D imaging stacks.

**Figure 2.**
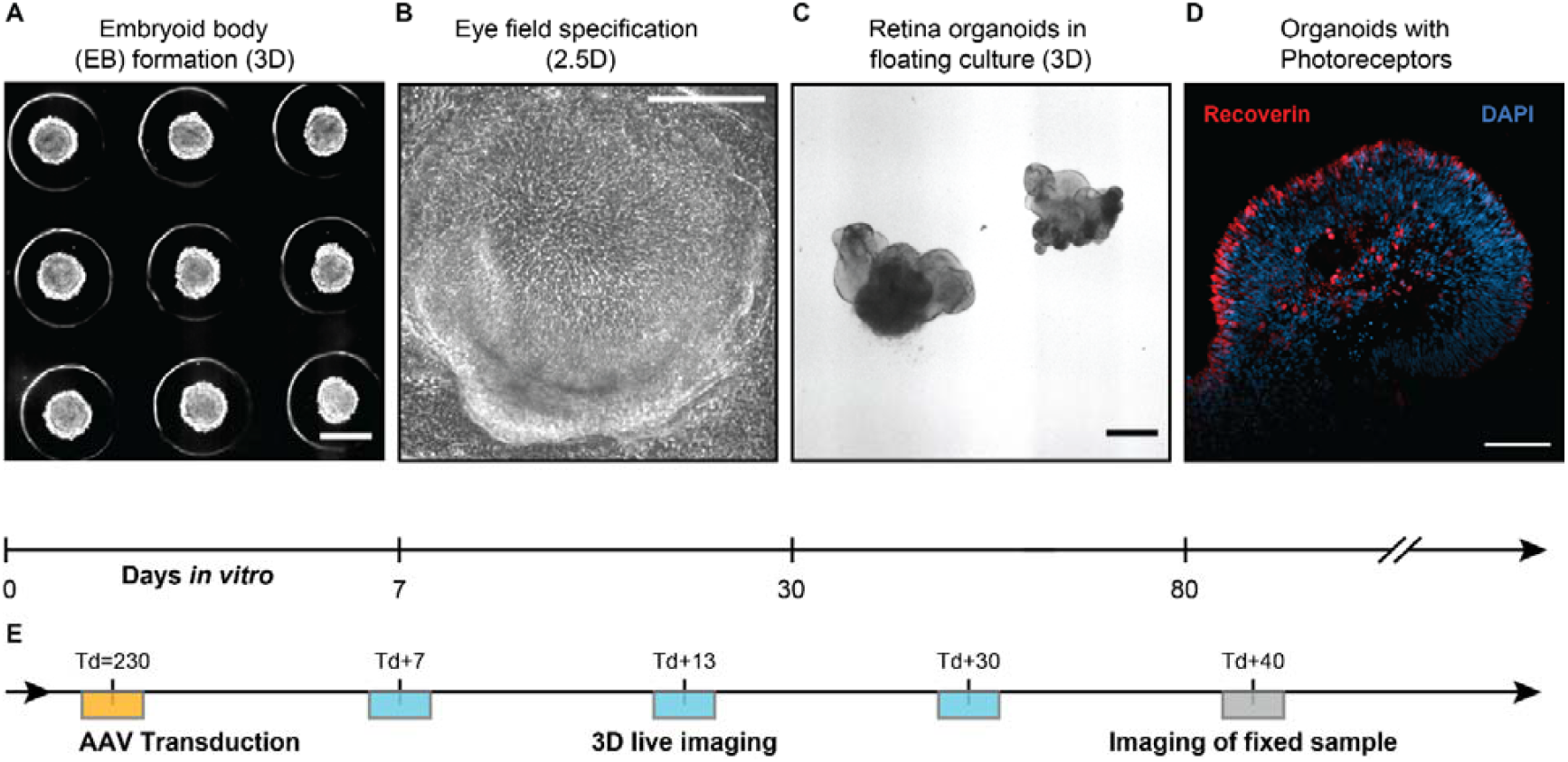
Human retinal organoids as a test bed for viral transduction. (A) Induced pluripotent stem cells (iPSCs) were transferred to 3D culture to form embryoid bodies (EB). (B) Re-plating the EBs onto 2D geometry (Matrigel matrix) for 21 days while supplying specific biochemical signals triggered the specification of the eye field (see Methods). (C) Floating culture enabled cell differentiation of retinal cell types. (D) 80 days old organoids displayed photoreceptor cells. (E) Transductions were carried out around day in vitro (DIV) 230 (yellow). Living transduced organoids were imaged and fixed 40 days after transduction (Td+40). Image types: (A-C) Brightfield, (D) immunofluorescence. Scale bars (A-C) 500 µm, (D): 100 µm.

### AAV transduction of human retinal organoids

To demonstrate and test our approach, we transduced human retinal organoids with three AAV vectors with different serotypes. In addition to using the clinically-tested serotype AAV2.7m8^[24]^ as a control, two optimized serotypes were selected for this study. Serotypes AAV2 and AAV9 have been previously reported to enhance the transfer of genetic material to retinal organoids.^[17]^ An engineered version of AAV2 (AAV2.NN) was used due to its superior performance compared to AAV2.7m8 with respect to transduction efficiency and onset kinetics.^[25]^ AAV9.NN is a modified AAV9 variant carrying the peptide insertion of AAV2.NN. Since potential clinical applications may benefit from testing in the highest maturity of human retinal tissue possible, we transduced hROs at a later stage of development on day in vitro (DIV) 230, marking day 0 after the day of transduction (Td+0) (Figure 2). Each organoid was supplied with 5*10^10^ vector genomes (see Methods). Eight hROs were transduced per serotype on consecutive days centered at Td = 230(−3, +4) days (Figure 2E, yellow box), imaged at three timepoints: Td+7, Td+13, Td+30, and fixed at Td+40 (Figure 2E, gray box).

### Confocal 3D live imaging

To obtain 3D images of the nuclei (Figure 3A) as well as of the AAV-induced eGFP fluorescence (Figure 3A), a commercial confocal microscope (Zeiss LSM980) was used in combination with a 40x NA (numerical aperture) 1.15 water immersion objective. The observation volume (211 × 211 × 100) µm^3^ was positioned in the sample such that the retinal cross-section filled the entire field of view at a depth of 100 µm. All measurements were taken using this standardized imaging volume. An experimental hurdle for imaging in deeper retinal layers is the wavelength-dependent penetration depth of the laser light, which can be increased e.g. using a 2-photon excitation source.^[26]^ We overcame this problem by using a near-infrared dye for the nuclei staining (excitation wavelength: 650nm, see Methods) and increasing the laser intensity with respect to imaging depth (see Methods). The illumination of both the nuclei and transduced channels was adjusted as a function of penetration depth while interpolating the intensity to z-depth containing fewer cells. Accordingly, the illumination intensity was adjusted to saturate one to two cells within each imaging plane. A 3D rendering of the raw data can be seen in Figure 3A (see also Supplementary Movie S1).

**Figure 3.**
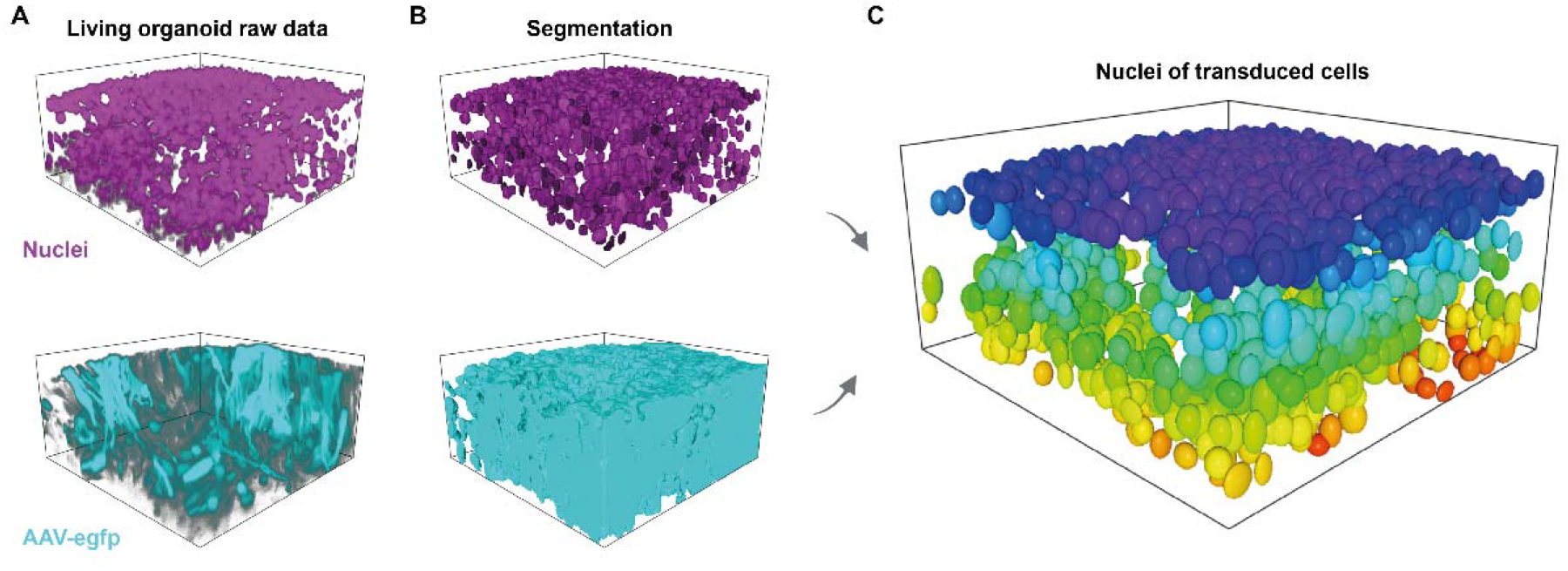
Quantification of transduction efficiency in living human retinal organoids. (**A**) Raw confocal image of 241-day old human retinal organoid transduced with AAV2.NN. (**B**) Segmentation of nuclei via deep learning^[21]^ and transduced volume via machine learning (see Methods). (**C**) Elliptical approximation of nuclei of transduced cells. Colour coding according to z-position. Observation volume b x w x h: 211×211×100 µm^3^.

### 3D Segmentation and quantification of transduction efficiency

The transduction efficiency is defined as *E* = *N*_*Td*_ / *N*_*Tot*_ where *N*_*Td*_ is the number of transduced cells and *N*_*Tot*_ the total cell number, respectively. Obtaining *N*_*Td*_ from 3D live confocal microscopy images of cytoplasmic reporters is challenging for human retinal organoids, but also for other neuronal tissues, since cells are densely packed, and substantially vary in aspect ratio and in fluorescence intensity due to cell type-specific transgene expression and transduction. To overcome this problem, we used a three-step approach: first, we segmented all nuclei using the deep-learning software, Cellpose (Version 2.0),^[21]^ in order to obtain *N*_*Tot*_, as well as the positions and minimal ellipsoidal representations of all nuclei (Figure 3B). This eliminated the requirement to distinguish single neurons in the harder-to-segment cytoplasmic channel.

In a second step, we segmented the transduced volume. In previous fluorescent-based approaches, the overall fluorescence was used as a proxy for the transduction efficiency.^[18]^ When transduction efficiency is evaluated this way, few transduced cells with a bright fluorescence signal and many transduced cells with low fluorescence intensity yield the same result (Figure 1B). To overcome this ambiguity, we trained a machine-learning algorithm on both low and high cytoplasmic intensity fluorescence signals (Figure 3B, see Experimental Section). The user-annotation is sensitive to a variety of image features like intensity gradients and shapes, enabling a more robust segmentation of the fluorescent volume compared to intensity-based thresholding alone. In order to adjust to the increasing illumination with imaging depth, we trained the segmentation algorithm using the transduced signal in both low-and high-intensity areas of the organoids at three depths (25, 50, and 75 µm). In a last step, we evaluated whether a neuron was transduced. Therefore, we calculated the overlap between the segmentation of the transduced volume with a thin virtual shell (thickness 0.8 µm) surrounding each segmented volume representing a nucleus.

In order to estimate the performance of our approach, we compared it to the manual counting method of assessing transduction efficiency. First, we selected three layers (at 25, 50 and 75 µm imaging depth) of different depths of a 3D stack. The cells in those layers were annotated manually by three different scientists as transduced or non-transduced based on the membrane stain and AAV-introduced GFP fluorescence (Figure 4A-D). Additionally, as a positive control (100%) we 3D-imaged endogenous eGFP expression, membrane, and nuclei stainings under control of the *Rax* (Rx) promotor in transgenic mouse retinal organoids (Figure 4E).^[26]^ As a negative control, untreated wild-type organoids were used (Figure 4F). After annotation, the manual counting results were then converted into a transduction efficiency by dividing the number of transduced cells by the total number of cells. Therefore, the mean transduction efficiency per layer, as well as the average efficiency and standard deviation over the means of all layers was calculated for human annotation. The results were then compared to the software application results of the same imaging stack. By trying different thresholds for the overlap between transduced volume and shell (Figure 4G, H), we chose the threshold (40%) resulting in the transduction efficiency closest to the manual upper and center layer results for our final threshold (Figure 4H).

**Figure 4.**
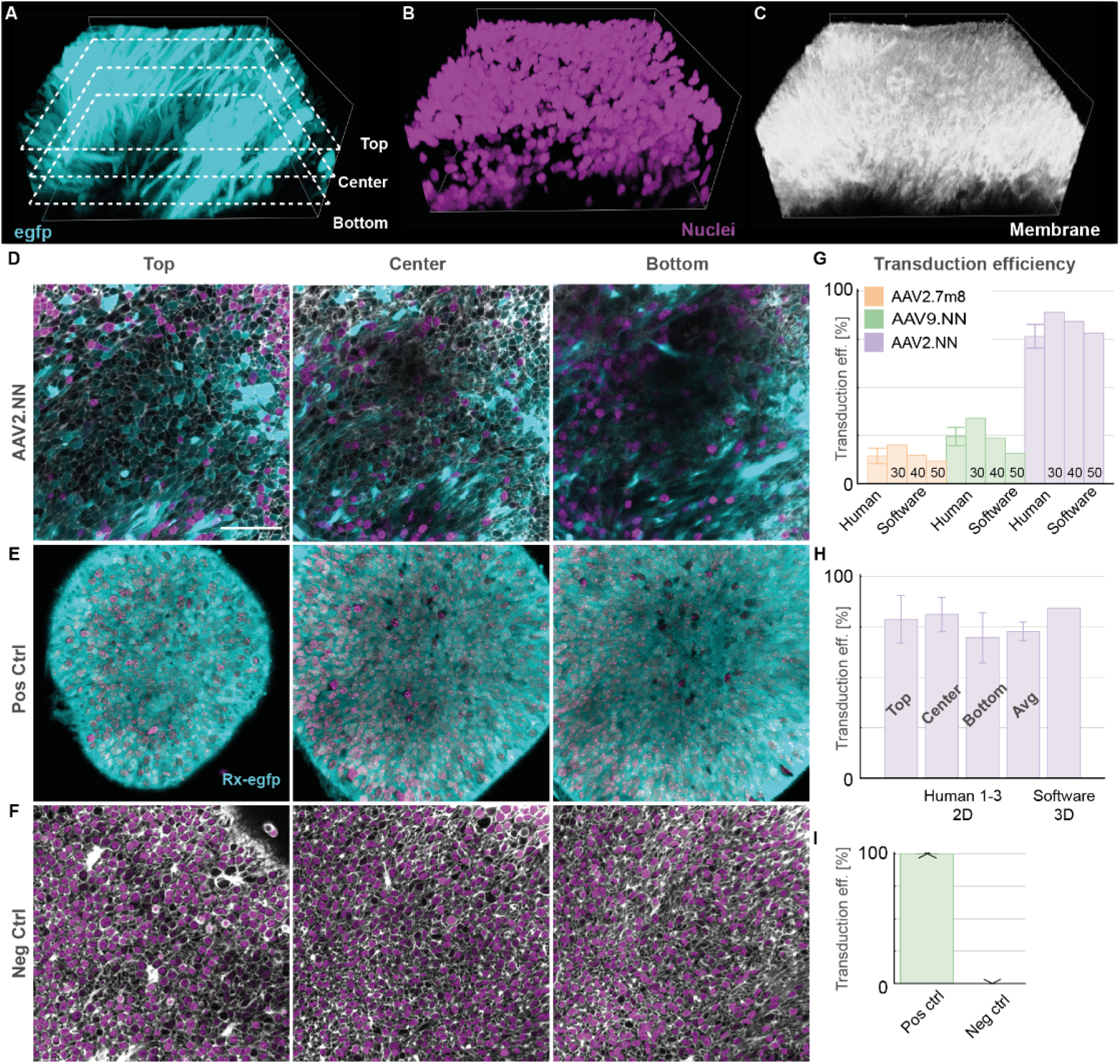
Assessment of the analysis pipeline and comparison with human counting. (A-C) 3D maximum intensity rendering of a confocal microscopy stack of a 239 days old human retinal organoid transduced with AAV2.NN (turquoise) stained for membrane (CellBrite 550; white) and nuclei (NucSpot 650; magenta). (D) 2D planes (top: ¼ of total depth (25 µm), center: ½ of total depth (50 µm), bottom: ¾ of total depth (75 of 100 µm)) from data (A-C) at different positions were used for manual assessment of transduced and total number of cells. (E) Positive control (Pos Ctrl) confocal images of the top, center and bottom plane of the 3D stack of an 18 days old mouse retinal organoid expressing eGFP under the Rx-promotor. (F) Negative control (Neg Ctrl) confocal images of the top, center and bottom plane of the 3D stack of a 264 days old human retinal organoid without virus. (G) Comparison of transduction efficiencies (eff.) as a function of shell volume threshold (see main text) and human annotation for three organoids transduced with different AAVs. Scale bar: 50 µm. Error bars indicate the standard deviation (SD; *N* = 3). (H) Transduction efficiencies of images in (D) each quantified via 3 human annotators. The efficiency for human annotation (3 annotators) averaged over the three planes (top, center, bottom) yields (73 ± 5)% (mean ± SD) compared to 84% for the software at a threshold of 40% shell to transduced volume overlap. See Supplementary Tables S2 and S3 for the counting and automated analysis data. (I) Quantification of transduction efficiency for positive (*N* = 3 different organoids) and negative (*N* = 8 different organoids) control stacks. Small error bars were made visible with an arrow.

For the automated analysis of the negative control, we obtained eGFP expression of 0% while the positive control showed eGFP expression in 99.9% of the cells (Figure 4I), highlighting the broad dynamic range of our approach.

We then used our pipeline to quantify the transduction efficiency in living hROs with three different AAV serotypes, AAV2.7m8, AAV2.NN and AAV9.NN, at three different time points (Figure 5). At a fourth timepoint, (Td+40), the organoids were fixed. Figure 5A shows the raw eGFP-signal of organoids transduced by the different vectors. The transduced nuclei’s ellipsoidal approximations are rendered in Figure 5B, and the mean transduction efficiency with standard error of the mean for all samples are plotted in Figure 5C.

**Figure 5.**
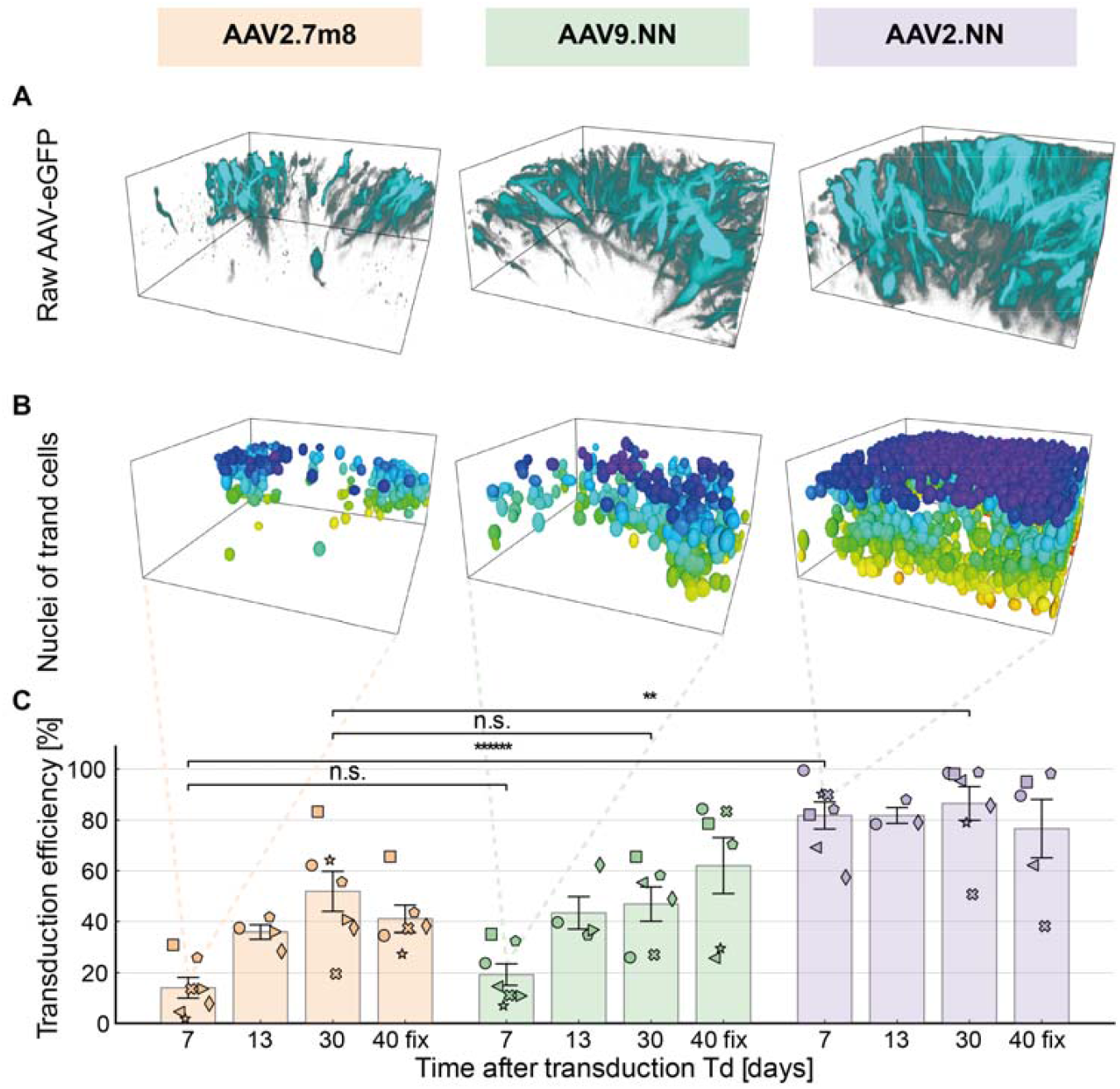
Transduction efficiency quantification for AAV serotypes AAV2.7m8, AAV2.9NN and AAV2.NN. (A) Representative raw images of living retina organoid 7 days after the day of transduction (Td). (B) Elliptical approximations of nuclei of transduced cells from (A). (C) Transduction efficiency over the span of 40 days after transduction. AAV2.NN has a faster onset and a higher overall efficiency compared to AAV2.7m8 and AAV9.NN. Each organoid is denoted with a different symbol (see Supplementary Tables S4-5). AAV2.7m8: *N* = (7,4,7,6); AAV9.NN: *N* = (7,4,6,6); AAV2.NN: *N* = (7,3,7,5) for (Td+7, Td+13, Td+30, Td+40). Td+7 until Td+30: living organoids; Td+40: organoids after fixation at day 40 were marked as “fix” (see Methods). Error bars indicate the standard error of the mean (SEM). Statistical significance: n.s.: *p* > 0.05, **: *p* ≤ 0.01, ******: *p* ≤ 10^−6^. Some 3D stacks were excluded from analysis when imaging purely retinal tissue was not possible.

The results obtained from living organoid samples (Td+30) agreed with the fixed samples (Td+40 fixed) within the error bars, indicating that fixed, whole-mount organoids yield comparable results to our live imaging approach. One week after transduction (Td+7) we observed a difference in efficiency between the tested serotypes AAV2.7m8, AAV9.NN and AAV2.NN (Figure 5A-C; Supplementary Tables S4-5). While serotypes AAV2.7m8 and AAV9.NN showed a similar onset time and overall level of eGFP expression, serotype AAV2.NN outperformed the other two variants substantially. Our work recovered the fast onset time of less than 7 days in transduction experiments performed with AAV2.NN^[17]^ as well as significant higher transduction efficiency found with AAV2.NN when compared to AAV2.7m8 with a 68% difference at 7 days post viral introduction.^[16]^ Most importantly, we can pinpoint the absolute amount of transduction efficiency, which reached a total (82 ± 5) % of the cells within the retinal organoid in the case of AAV2.NN (Td+7) (Figure 5C).

### Analyzing transduction efficiencies of different areas of depth within 3D image stacks

To demonstrate the use of our method in estimating the transduction efficiency of certain regions within our samples, e.g. the outer nuclear layer (ONL) consisting of primarily photoreceptors, we assessed the transduction efficiency at different imaging depths (Figure 6A, Supplementary Figure S3). Therefore, we measured the depth of the border between the ONL and the inner nuclear layer (INL), visible through a gap in nuclear density located at about 40 µm from the first imaging plane.

**Figure 6.**
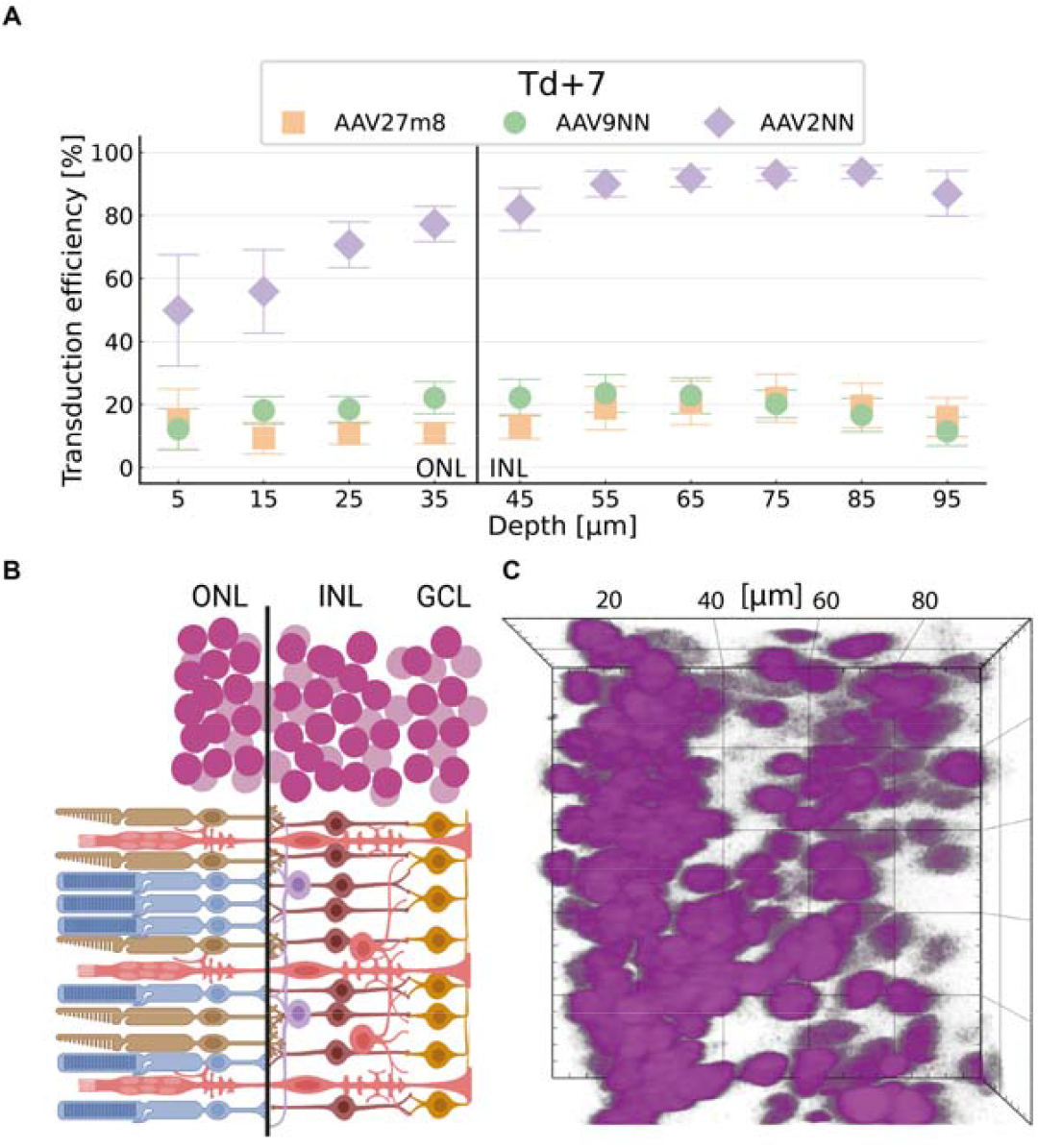
Spatially resolved quantification of viral vector transduction efficiency in living retinal organoids. (A) The normalized transduction efficiency was determined at Td+7 (See Supplementary Figure S3 for all timepoints). The position of retinal neurons was determined by the nuclei centroid position. The transduction efficiency was then binned over 10 µm (z-axis) deep slices across the 3D stack (see Methods, Supplementary Table S6) and plotted against the imaging depth (µm) for the three different viral serotypes; AAV2.NN shows high levels of transduction in deeper layers already on Td+7. AAV2.7m8 and AAV9.NN show significant differences in transfection at different depths; sample size (number of organoids) for each serotype: *N* = 7. Error bars indicate the standard error of the mean (SEM). The black solid line depicts the estimated border between the outer nuclear layer (ONL) and inner nuclear layer (INL) illustrated in (B): Sketch of the retinal layers ONL consisting of primarily photoreceptors (rods and cones), INL with nuclei of bipolar cells, horizontal cells, amacrine cells and Müller glia cells as well as the ganglion cell layer (GCL). (C) Example of nuclei layering in a 100 × 100 × 40 µm section of a retinal organoid 3D stack taken at Td+7. The ONL was separated visually from the INL with a black line at 40 µm of imaging depth (z-axis). One square in the image measures 20 × 20 µm^2^.

We found that, on average, when AAV2.NN was applied, cells of the ONL showed a higher transduction efficiency than the cells of the INL 7 days after transduction (Figure 6A). The other two implemented serotypes, AAV2.7m8 and AAV9.NN, showed no statistically significant differences for this timepoint (Figure 6A). Further, these data show that not only does AAV2.NN have a faster transduction time in general, but its efficiency at depths greater than 50 µm is more than four-fold as high as that of the other vectors.

This demonstrates that our technique enables us to observe the spatial distribution of transduction over time throughout different parts of the retina (Figure 6). In addition, we documented the organoid growth over the 40-days imaging span with brightfield imaging (Supplementary Figure S4-6).

## Discussion

We have developed a pipeline for quantitative assessment of drug delivery in intact living human organoids, providing a versatile tool to the community. Traditional methods to assess gene delivery using fluorescent reporters either failed to provide spatial information, such as flow cytometry,^[18]^ or require tissue fixation, which prevents dynamic studies.^[17,22]^ We combined confocal microscopy, recent segmentation tools and a robust quantification algorithm and generated a platform which provides transduction efficiency spatially and temporally resolved in 3D. To uniquely assess whether a neuron is transduced or not, we considered the segmentation of the fluorescence signal within a thin shell around each nucleus. To detect transduced cells in the presence of background signal, we trained a machine-learning model instead of assessing the fluorescence signal directly. This allowed us to assess the transduction of thousands of cells per volume with a performance comparable to human annotation. We used our platform to quantify AAV transduction efficiency of three different serotypes. We found that AAV2.NN outperforms the serotypes AAV9.NN and AAV2.7m8 with respect to onset time (< 7 days), overall transduction efficiency (82%) and the efficiency to transduce deeper retina layers.

The ability to temporally follow individual organoids and assess their transduction at a single neuron level provides two key benefits for the community. First, it allows optimizing AAV serotype selection and AAV delivery protocols with respect to their onset time, thus minimizing the time for a gene therapy to become effective in the body. Second, the impact of the AAV on the organoid can be tracked and safety margins can be effectively established.

Aside from the methodological benefits, our approach brings spatiotemporal quantification to organoid research laboratories due to the comparably low technological hurdles. The application requires a standard confocal microscope to image the fluorescent reporters of the gene delivery system to be examined, as well as commonly used nuclear live stains. Our analysis approach is simple yet robust with only one key parameter, leveraging the segmentation power of recent deep learning tools. Lastly, time-consuming cryo-sectioning as typically done can be avoided by allowing longitudinal live imaging of the same organoids.

The technology introduced here can be readily used in other organoid systems, both neuronal such as brain organoids and non-neuronal. It could be expanded to organ explants, including retina explants and brain slices and any other 3D cell culture system. While the approach has been optimized to work with densely packed tissue, adaptation of the nuclei shell overlap threshold allows assessment in tissues with different cell densities. Our approach will guide the optimization of gene delivery to specific cells when used in conjuncture with cell type-specific reporter lines such as GFP-labelled retinal ganglion cells.^[27]^ This could provide deeper insight into AAV serotype selectivity.^[28]^

Our platform has the potential to leverage the efficiency and specificity of the quantification of further transfection and transduction techniques in a spatiotemporal way, such as mRNA delivery^[15]^ and nanoparticles^[29]^ and lentiviral transduction. On a broader scope, our method has the potential to improve the development of therapies for various tissues beyond the retina.

## Methods

### Statistical analysis

#### Pre-processing

‐ Figure 4G-H: Representative samples were chosen based on membrane staining
‐ Figure 4I: Contains all samples
‐ Figure 5C, Figure 6A, Supplementary Figure S3: Images were excluded from data evaluation if there was tissue other than retinal tissue growing in the imaged volume.

#### Data presentation

All errors are given as standard error of the mean, 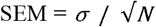, with the unbiased sample standard deviation (SD), 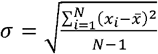, the sample size *N* and the sample’s mean 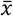.

‐ Figure 4G: mean ± SD
‐ Figure 4H: mean ± SD
‐ Figure 4I: mean ± SD
‐ Figure 5C: mean ± SEM
‐ Figure 6A: mean ± SEM
‐ Supplementary Figure S3: mean ± SEM

#### Sample size

‐ Figure 4G: *N*_*layers*_ = 3
‐ Figure 4H: *N*_*layers*_ = 3, *N*_*annotators*_ = 3
‐ Figure 4I: *N*_*pos*_ = 3, *N*_*pos*_ = 8
‐ Figure 5C, Figure 6A, Supplementary Figure S3: AAV2.7m8: *N* = (7,4,7,6); AAV9.NN: *N* = (7,4,6,6); AAV2.NN: *N* = (7,3,7,5) for (Td+7, Td+13, Td+30, Td+40) (See Supplementary Table S2 for the number of areas per organoid).

#### Statistical methods

Figure 5C: First we tested for equal population variance by performing a Levene’s test using the group median and α = 0.05. Because all data sets passed the test, we assumed an equal population variance and then performed an unpaired t-test with α = 0.05 to obtain the p-values reported in Figure 5C.

#### Software

Statistical evaluation was performed in a custom Python code. The functions sqrt, mean and std from the package numpy were used for calculating the mean, the standard deviation and the standard error of the mean. The functions ttest_ind and levene from the package scipy.stats were used for evaluating the statistical significance.

### Human iPSC culture

The commercially available cell line iPS(IMR90)-4 (WiCell®, ^[30]^) was cultured according to WiCell® protocol. Matrigel was prepared by dissolving Matrigel (1 mg, Corning® Matrigel® Basement Membrane Matrix Growth Factor Reduced, 354230, lot number: 0295001) in cold DMEM/F-12 medium (1 ml, Glutamax, GIBCO, 31331-028). The dissolved Matrigel solution was further diluted with the addition of 11 ml cold DMEM/F-12 medium. 1 ml of diluted Matrigel solution was plated per well in a 6-well plate which was then incubated at 37 °C for 1-2 hours. Once the Matrigel set, DMEM/F-12 medium (1 ml) was added to each well to prevent drying. Plates were kept at 37 °C overnight. IPSCs were thawed in a 37 °C water bath and subsequently diluted with mTeSR1 medium (Stem Cell Technologies, 85850). The cells were centrifuged at 200 g for 5 minutes, supernatant was discarded, and the cells were resuspended in mTeSR1 medium (0.5 ml/well to be seeded) supplemented with Rock Inhibitor (RI) (10µM, Stem Cell Technologies, Y-27632) for every 6-well that should receive cells. Medium removed from the prepared Matrigel coated wells and 0.5 ml of cells were plated per well. The iPSCs were moved to the incubator (37 °C) and were fed with 2 ml culture medium daily. Once the colonies became too dense or increased differentiation occurred, the iPSCs were passaged.

Passaging (1:2 ratio) was conducted using EDTA (Sigma-Aldrich, E8008) as described in the WiCell® protocol. Areas containing differentiated cells were removed manually using a pipette tip under a stereomicroscope. Media was aspirated from each well and wells were subsequently rinsed with PBS (1 ml, Dulbecco, Biochrom, L182-50). PBS was then aspirated from each well. To remove the iPSCs from the surface of the well, 1 ml EDTA was added at RT (room temperature) and incubated for 6-9 minutes. Meanwhile, new culturing dishes were prepared by replacing the medium in new Matrigel coated wells with 1 ml of mTeSR1 medium per well. Following the incubation period, EDTA was aspirated and the cells were detached gently from the surface using 2 ml culture medium and a pasteur pipette. 1 ml of homogenous cell colonies were plated onto each new Matrigel coated well, distributed evenly by slight movement of the dish, and incubated overnight.

### Human retinal organoid culture

hROs were generated in agarose microwell arrays which restrict the size of growing organoids following the protocol described in Cowan et al with minor alterations. This method has been found to efficiently generate organoids while requiring less iPSCs than alternative approaches.^[7]^

Agarose molds were prepared for iPSC culture. MicroTissues® 3D Petri Dish® micro-mold spheroids (Sigma, Z784019) were filled with agarose (2%, Thermo Fisher Scientific, R0491) dissolved in DMEM (Thermo Fisher Scientific, 10569-010). The solidified agarose molds were transferred into a 12-well dish. The molds were equilibrated by adding mTeSR1 medium (1.5 ml) was added to each well and the plates were incubated at 37 °C for a minimum of 15 minutes. This equilibration step was repeated with fresh mTeSR1 medium. The molds were stored at 4 °C and a final equilibration was done with fresh mTeSR1 media.

Before seeding iPSCs into the microwells, the cells were plated and passaged once in a Matrigel coated 6-well plate as described above. To dissociate the iPSCs from the 6-well plate, each well was washed twice with PBS (1 ml) and incubated for 10 minutes in PBS (600 µl) with EDTA (0.5 M) at room temperature. After aspirating the PBS containing EDTA, the plate was incubated in Accutase (500 µl, Invitrogen, 00-4555-56) for 3 minutes at 37 °C. mTeSR1 (1 ml) with RI (10 µM) was added and the cells were pipetted up and down to generate a single-cell suspension. Cells were pelleted for 5 minutes at 200g, supernatant was discarded, and the cells were resuspended in mTeSR1 (5 ml) with RI (10 µM). To seed, the cells were diluted to achieve a concentration of approximately 60,000 cells in 150 µl and 150 µl of cell suspension were seeded per agarose well. To allow the cells to settle, the plate was placed in the incubator (37 °C) for 30 minutes, after which the well was filled up with mTeSR1 (1.5 ml) with RI 10 (µM). The plates were kept in the incubator at 37 °C and fed according to the following.

On day 1, a third of the medium was replaced with NIM medium containing DMEM/F12, N2 supplement (GIBCO, 17502-048), non-essential amino acid (NEAA) solution (1%, SIGMA, M7145), and Heparin (2 µg/ml, SIGMA, H3149). On day 2, half of the medium was replaced with NIM medium and on day 3, the entire medium was replaced with NIM medium. From day 4 to day 6, 1.5 ml NIM medium were added daily.

On day 7, the embryoid bodies were transferred to a Matrigel coated 6-well plate and the NIM medium was replaced daily.

From day 16 on, NIM medium was replaced with 3:1 medium containing DMEM (72%, GIBCO, 10569-010), F12 (24%, GIBCO, 31765-027), B27 w/o VitA (2%, GIBCO, 12587-010), NEAA (1%, Sigma, M7145), and Pen/Strep (1%, GIBCO, 15140-122). The medium was replaced daily.

On day 31, retinal structures were detached by checkerboard scraping scratching a 1-2 mm grid with a 200 µl pipette tip. This causes retina structures to come off in larger tissue pieces. Debris and single cells were removed by washing the aggregates three times in a 15 ml tube in 3:1 medium by sedimentation. The aggregates were maintained in suspension in 6-7 ml 3:1 medium in 6 cm plates. The media of the floating cultures was changed three times per week.

One week after checkerboard scraping on day 38, aggregates that did not contain phase-bright stratified neuroepithelium were manually removed.

From day 42, the 3:1 medium was supplemented with heat-inactivated FBS (10%, Sigma-Aldrich, ES009-B) and taurine (100 µM, T0625-25G, dissolved in MilliQ H2O). Organoids were fed three times a week.

After day 70, the aforementioned 3:1 medium supplemented with FBS and taurine was further supplemented with retinoic acid (1 µM, Sigma, R2625, dissolved in DMSO).

From day 98 on, the B27 supplement in the 3:1 medium was replaced with N2 supplement (1%, GIBCO, 17502-001) and retinoic acid was reduced to 0.5 µM.

#### Fixation

Retina organoids were fixed in PFA (4%, Sigma-Aldrich, F8775-25ML) in PBS overnight at 4 °C. Afterwards, the organoids were cryopreserved in PBS containing sucrose (30%, Sigma, S1888-500G) overnight at 4 °C. For the fixed TP4 measurements, organoids where incubated for 48 hours in PBS, TritonX-100 (0.5%, Roth, 3051.3) and stained with DAPI and 1:500 WGA-CF (Supplementary Table S1).

#### Cryosectioning

2-4 organoids were embedded in Neg-50™ Frozen Section Medium (Thermo Fisher Scientific, 6502B) and frozen on dry ice. Cryosections (10 µm) were cut using a Epredia™ CryoStar™ NX70 cryostat. The slices were picked up directly with SuperFrost Plus™ Adhesion slides (Epredia™ J1800AMNZ). The slides were left at RT to dry for at least 2 h and stored at -80 °C.

#### Staining

Slides were left at RT to thaw and dry for 30 minutes and rehydrated in PBS until the embedding medium was dissolved and only the organoids remained on the slides. The sections were incubated in blocking solution containing PBS, Triton X-100 (0.5%, Roth, 3051.3), and BSA (1%, Sigma, A7638-10G) for 1h at RT.

The blocking solution was aspirated and primary antibodies (Supplementary Table S1) diluted in blocking solution were added to the slides and incubated overnight at 4 °C. Slides were rinsed in PBS for ten minutes twice. Secondary antibodies diluted in blocking solution, DAPI, and the membrane stain WGA CF555 were added to the slices and incubated for 2 hours at RT. Slides were rinsed in PBS for ten minutes twice and subsequently rinsed once more in distilled water. Coverslips were mounted to the slides using Mounting Medium Fluoromount-G™ (Thermo Fisher Scientific, 00-4958-02).

### AAV vector production

AAV vector production was performed as previously described.^[25,31]^ A self-complementary AAV plasmid containing a cytomegalovirus-enhanced green fluorescent protein (CMV-eGFP) expression cassette flanked by ITRs from AAV2 ^[32]^ was used as the vector genome plasmid.

### Transduction experiments

Per condition, eight organoids were transduced, with the first transduction (T1) taking place at DIV227. The procedure was repeated daily until DIV 235 with one organoid per condition per day. At the day of transduction (Td), 5*10^10^ viral genomes/organoid were added, and the viral solution was filled up to the final volume (300 µl) with culture media. On Td+1 (one day after Td) the organoids were transferred into the wells of a 24-well plate. Each well was filled up to 1 ml with medium. On Td+4, the organoid was washed twice with PBS and supplied with 1 ml fresh media. Then the organoids were integrated into the regular feeding schedule of three weekly exchanges of half the volume with fresh media. After imaging of Td+30 the media was replaced completely with fresh media.

### Organoid staining for live imaging

Nuclei were stained using Nuncspot Live 650 (Biotium, 1:250). Membranes were fluorescently labeled using CellBrite Steady 550 (Biotium, 1:1000). To this end, organoids in were placed in glass-bottom dishes, the stains were added to the medium and organoids were incubated at 37 °C for 2 hours. Organoids were imaged using a confocal microscope (LSM980, Zeiss) in combination with a water immersion objective (40x). The membrane channel was used as a guide to identify the retinal tissue within the organoid which was then exclusively imaged. Volumes which contained other tissue types such as RPE (retinal pigment epithelium) were discarded. A sampling of (1024×1024×400) voxel was chosen corresponding to an observation volume of (211×211×100) µm^3^.

### Analysis pipeline

In order to extract transduction efficiency information from the confocal 3D images, both the segmentation of the nuclei channel and the cytoplasmic eGFP expressing cells was obtained. The segmentation was performed using the Arivis Vision4D (Zeiss) software package in several steps:

1. Conversion of the raw images (1024×1024×400) voxel into the arivis .sis format.
2. Denoising of the channels using the Denoise option
3. Nuclei segmentation via Cellpose ^[21]^ interfaced via python. For the cellpose segmentation following settings were used:

a. Binning to 50%
b. Discrete Gaussian denoising (0.829 µm diameter)
c. Cellpose custom model: CPx (Cellpose 2.0)
d. Thresholds: estimated diameter in µm: 5.0; flow threshold: 1.0, Mask inclusion threshold (cellprob. threshold): -6; maximal tile size: 4096 pixels, nuclei object volume filter [10 µm^3^, 1 m^3^]
4. Segmentation of the transduced volume (cytoplasmic eGFP) via the Arivis Machine Learning trainer. Settings were:

a. Arivis Machine Learning probability training for segmentation (trained on microscopy images of organoids to recognize areas with fluorescence signal as transduced and the background as not transduced)
b. Intensity threshold segmentation: Set range to exclude probabilities lower than 50%
c. Set object threshold: Only objects larger than 10 µm^3^
5. Identification of nuclei of transduced cells (nuclei that are located within a transduced environment):

a. Set Nuclei size threshold to larger than 100 µm^3^
b. Dilated nuclei objects by 2 pixels
c. To obtain the virtual shell around the nuclei subtracted intersection calculation (A-B) between the dilated nuclei objects (A) and the non-dilated nuclei objects (B)
d. Removed artifacts by applying a threshold of 50 µm^3^ (human organoid tissue; 30 µm^3^ for mouse organoid tissue) minimal size to the shells (determinable by the ratio between the number of nuclei to the number of shells (≈ 1))
e. Calculated compartments of the transduced volume segmentation to the size filtered shells (overlap threshold > 40%). The threshold is determined as shown in Figure 4G-H.
f. To obtain nuclei of transduced cells: Filter shells by overlap ≥ 0.4 (depending on threshold chosen for step e.)
g. Export summary for the total nuclei count (size filtered shells) and the number of transduced nuclei (shells with overlap ≥ 0.4)
h. Obtain transduction efficiency by dividing the sum of shells with overlap ≥ 0.4 by the sum of size filtered shells

### Comparison with human counting

One confocal image stack taken at Td+7 was selected for each AAV serotype. Three planes were extracted from each stack at depths of 25 µm, 50 µm and 75 µm (total stack depth: 100 µm, Figure 4A-D). The total cell number and the number of transduced cells were then independently counted by three humans using the AAV2.NN sample. The manual counting was based on the membrane label and the fluorescence signal from the AAV-delivered eGFP and performed using ImageJ. Both cells with low and high fluorescence were counted as transduced. Based on the comparison of the average human result with the pipeline, a suitable shell threshold for the pipeline was established (40%). In a final step, the selected threshold was tested by comparing the result of two other image sets (AAV2.7m8, AAV9.NN) with human annotation.

### Transduction efficiency quantification

The transduction efficiency *E* was calculated by dividing the number of transduced cells in a given organoid or volume (*N*_*Td*_) by the number of segmented cells in the same volume (*N*_*Tot*_), *E* = *N*_*Td*_ / *N*_*Tot*_. If there were measurements of several areas of the same organoid, then the transduction efficiencies of all areas were averaged and the obtained mean was treated as if it was a single measurement of an organoid. Then, the mean transduction efficiency at a certain day was calculated by averaging over the transduction efficiencies of all organoids at that day (Figure 5C, Supplementary Tables S4-5).

For determining the depth profile of transfection, all segmented cells in layers of thickness 10 µm were first binned for each organoid. The binning is necessary because no segmented cell is located at exactly the same depth. Next, the transduction efficiency was calculated separately for each bin. Third, the transduction efficiencies of the bins at certain depths (e.g. bin “5 µm” ranges from 0 µm to 10 µm) of different areas in one organoid were averaged and the obtained mean was treated as if it was a single measurement of an organoid. Finally, the transduction efficiency of the bins was averaged over all organoids. If there were no segmented cells present in a bin then this bin was excluded from averaging (Figure 6A, Supplementary Figure S3, Supplementary Table S6).

## Data availability

### Lead contact

Further information and requests for resources and reagents should be directed to and will be fulfilled by the lead contact, Friedhelm Serwane.

### Materials availability statement

- No plasmids were generated in this study.

### Data and code availability

- Microscopy data reported in this paper will be shared by the lead contact upon request.
- Any additional information required to reanalyze the data reported in this paper is available from the lead contact upon request.

## Acknowledgements

We thank Sebastian Willenberg and Guillermo Prol Castello for their contributions in the segmentation software. This work was supported by the European Research Council (ERC) under the European Union’s Horizon 2020 research and innovation program (Grant Agreement No. 850691), the Center for NanoScience (CeNS) at the Ludwig-Maximilian-University (LMU) Munich and the Munich Cluster for Systems Neurology (SyNergy). In addition, it was supported by the Deutsche Forschungsgemeinschaft (DFG, German Research Foundation) – Forschergruppe Project number 513025799. Figure 1, Figure 6B and the Table of Content illustration were created in BioRender. Rogler, T. (2025) https://biorender.com/ztu0mn5; https://biorender.com/o6jx5gz.

## Author contributions

S.M., F.S. and K.S. conceived the research. K.S., S.S., A.T. and R.R. cultured the human organoids, R.J. and T.S.R. the mouse organoids. K.S., T.S.R. and F.S. developed the analysis pipeline. T.S.R. and A.B. analyzed the data. S.B. and S.M. prepared the AAVs. K.S. and A.T. performed transduction experiments. Confocal imaging was performed by K.S., S.S. and A.T. (human organoids) and R.J. and T.S.R. (mouse organoids). K.S. and S.S. performed the immunohistochemistry. A.B. and E.S. performed the statistical analysis. All authors contributed to the manuscript.

