## Supplementary Information for "3D Quantification of Viral Transduction Efficiency in Living Human Retinal Organoids"

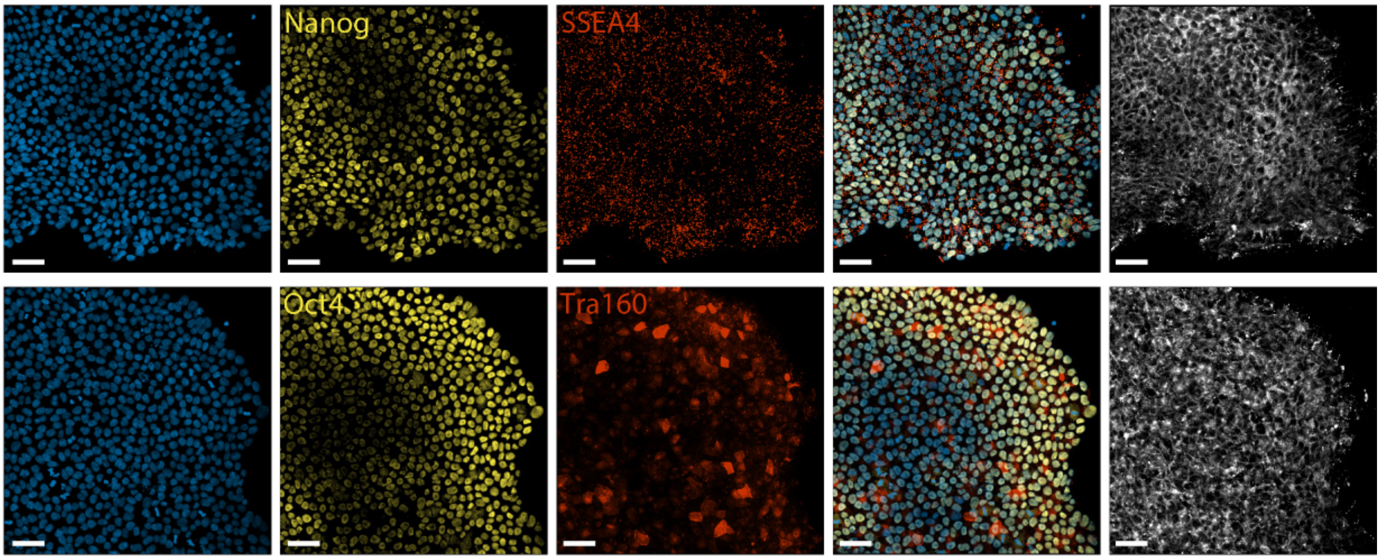


**Supplementary Figure S1.** Confocal images of pluripotency staining of iPS(IMR90)-4p37. Primary antibodies indicated in yellow and red, blue: DAPI, white: WGA-CF membrane stain. Scale: 50 microns.

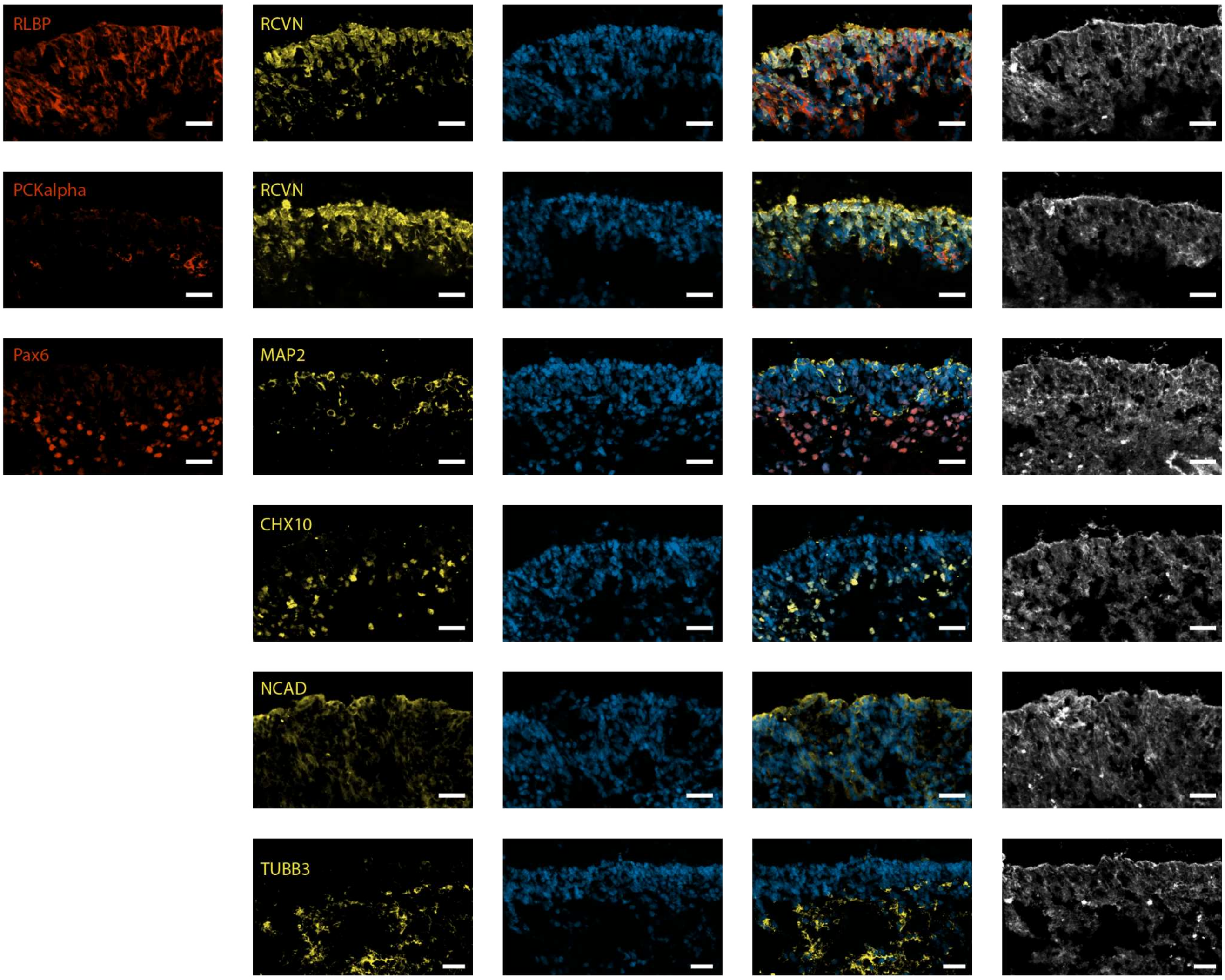


**Supplementary Figure S2.** Confocal images of immunofluorescence staining DIV227 human retinal organoids. Primary antibodies indicated in red and yellow, blue: DAPI, white: WGA-CF membrane stain. Scale bar: 30 microns. The markers were selected to highlight retina typical markers: RLBP (Retinaldehyde binding protein 1; müller glia cells), RCVN (recoverin; photoreceptor marker), PKCalpha (protein kinase C alpha; rod bipolar cells), Pax6 (paired box 6; neuronal progenitors), Map2 (microtubule associated protein 2; neuronal marker), CHX10 (visual system homeobox 2; retinal progenitor cells, bipolar cells), NCAD (N-cadherin; cell-cell adhesion, role in retina lamination) , TUBB3 (tubulin beta 3; microtubule, axone)

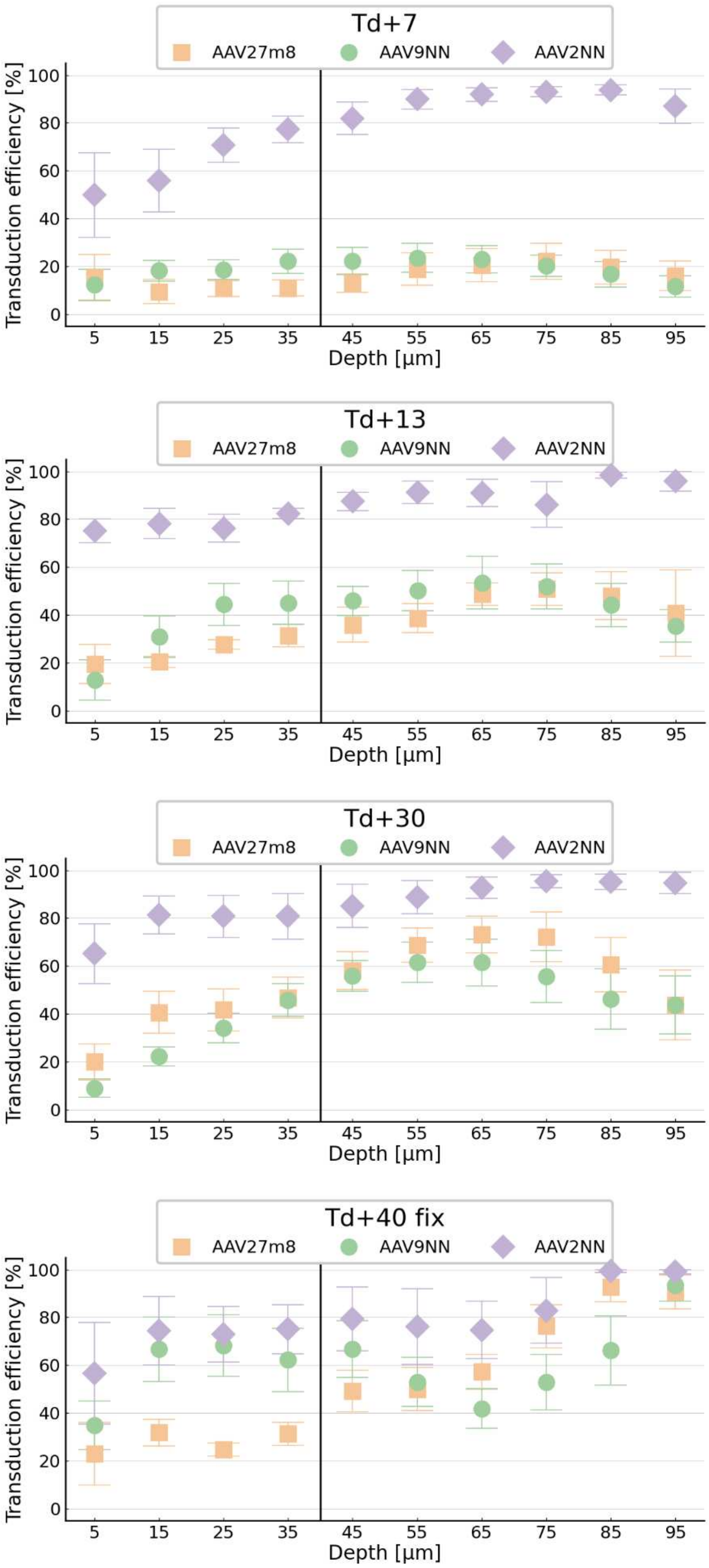


**Supplementary Figure S3.** Additional data for Figure 6A. Spatially resolved quantification of viral vector transduction efficiency in living retinal organoids at all timepoints. Transductions efficiency was binned over 10 µm deep z-slices. Error bars indicate the standard error of the mean (SEM).

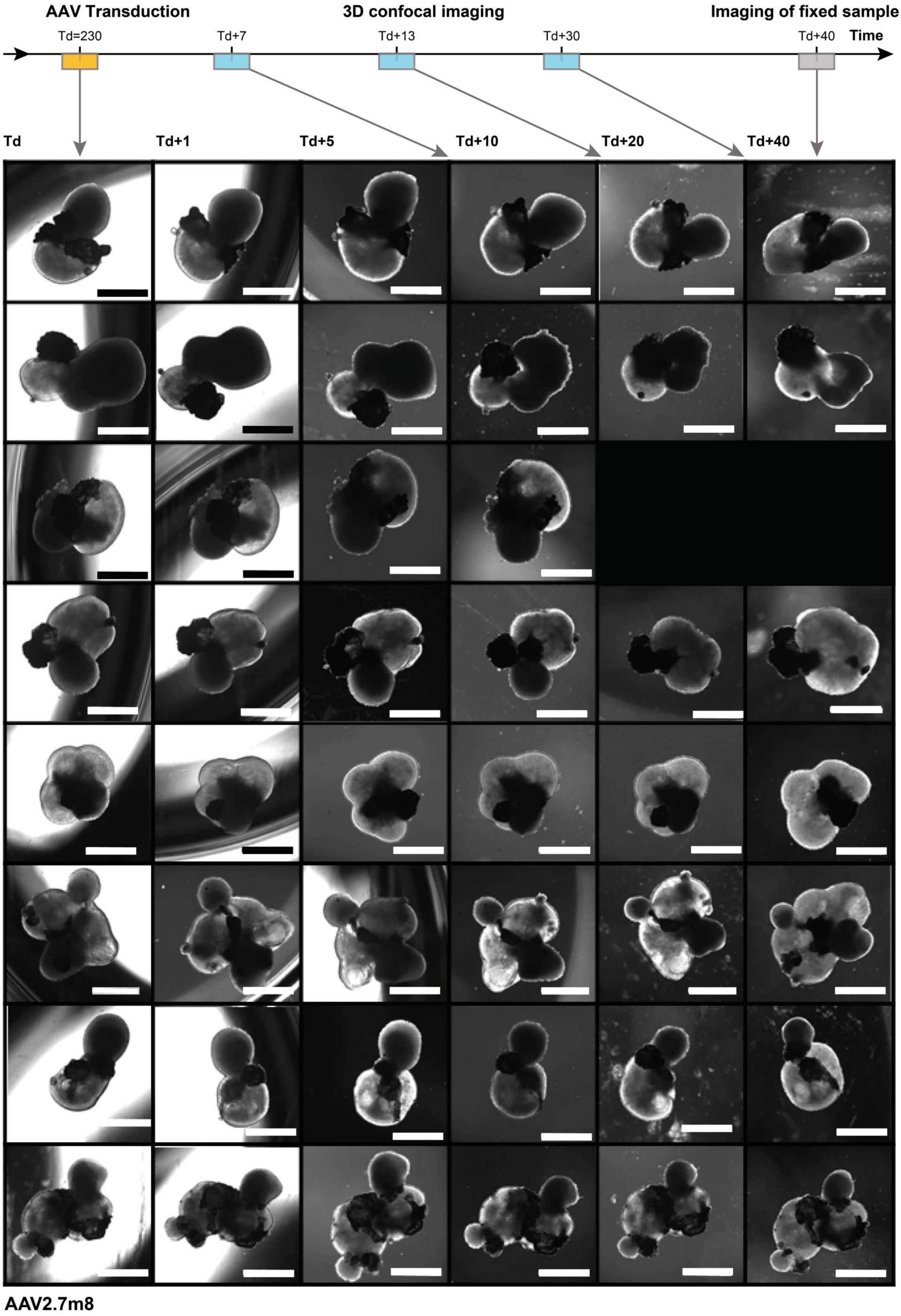


**Supplementary Figure S4.** Brightfield images of 8 human retinal organoids transduced with AAV2.7m8 and imaged at six time points after transduction (Td), related to Figure 4. Organoids from transduction experiment T1-8 from top to bottom. T1 was transduced at DIV227, T8 at DIV235. Scale bars: 500 microns.

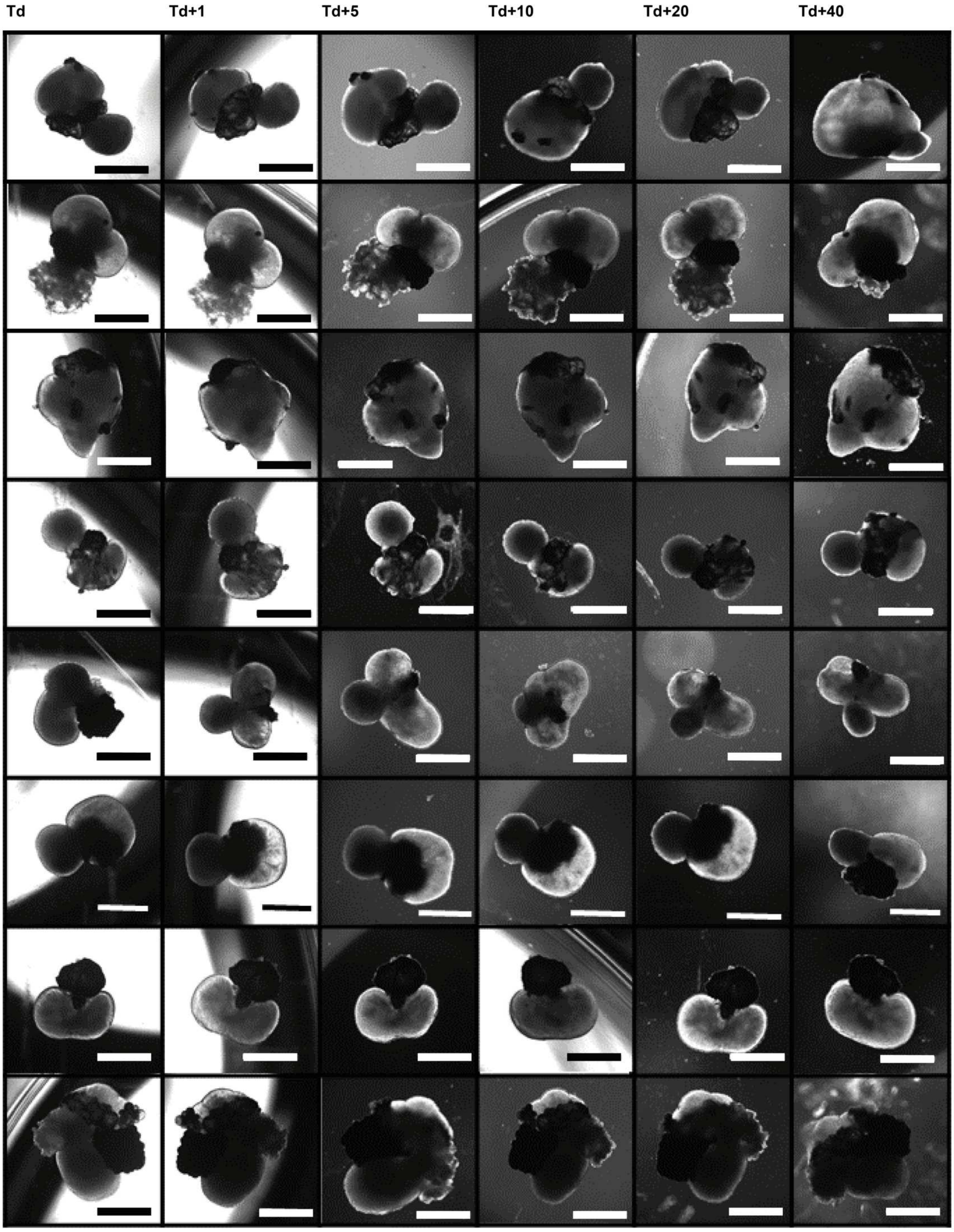


**Supplementary Figure S5.** Brightfield images of 8 human retinal organoids transduced with AAV9.NN and imaged at six time points after transduction (Td), related to Figure 4. Organoids from transduction experiment T1-8 from top to bottom. T1 was transduced at DIV227, T8 at DIV235. Scale bars: 500 microns.

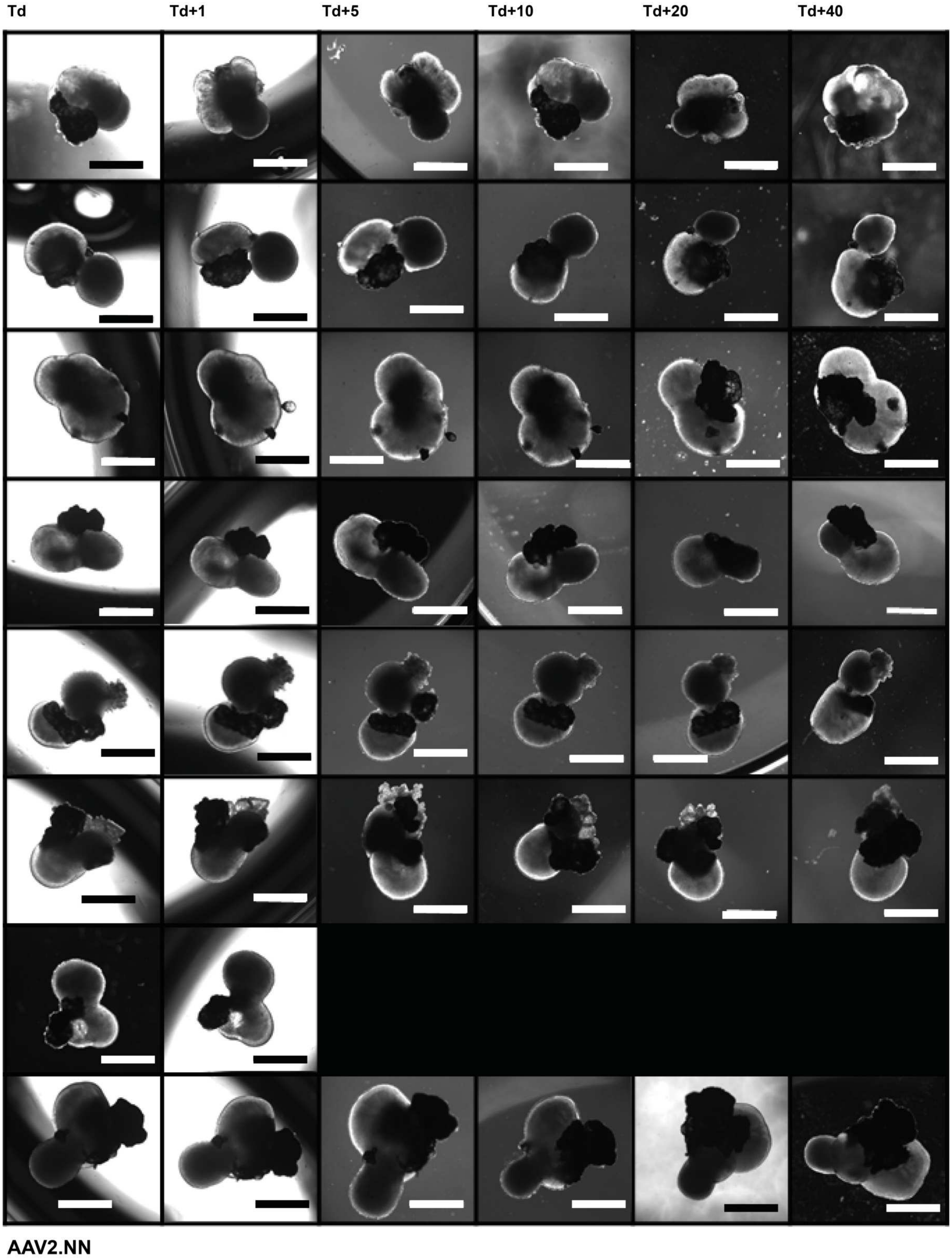


**Supplementary Figure S6.** Brightfield images of 8 human retinal organoids transduced with AAV2.NN and imaged at six time points after transduction (Td), related to Figure 4. Organoids from transduction experiment T1-8 from top to bottom. T1 was transduced at DIV227, T8 at DIV235. Scale bars: 500 microns.

**Supplementary Table S1.** Reagents used for live imaging, staining of cryosectioned organoids, and pluripotency staining of iPSCs.

| Antibody | Manufacturer | Product number | Dilution (Concentration) |
|---|---|---|---|
| Recoverin | Sigma | AB5585 | 500 |
| RLBP1/CRALBP | abcam | AB15051 | 100 |
| RBPMS | PhosphoSolutions | 1830-RBPMS | 100 |
| RBPMS-thermo | Thermo Fisher Scientific | PA5-31231 | 100 |
| Tuj1 | Biolegend | 801213 | 100 |
| PCKalpha | Santa Cruz Biotechnology | sc-8393 | 100 |
| n-Cadherin | Thermo Fisher Scientific | PA5-29570 | 200 |
| Rx | Santa Cruz Biotechnology | sc-271889 | 200 |
| MAP2 aves | Aves labs | MAP-0020 | 500 |
| Pax6 | GeneTex | GTX113241 | 100 |
| ZO-1 | Thermo Fisher Scientific | 40-2200 | 100 |
| CHX10 | Thermo Fisher Scientific | PA1-12565 | 200 |
| Nanog | Gift from D. Paquet | | 500 |
| Oct4 | Gift from D. Paquet | | 500 |
| SSEA4 | Gift from D. Paquet | | 100 |
| Tra160 | Gift from D. Paquet | | 500 |
| DAPI | Invitrogen | D1306 | (0.1 µg/ml) |
| WGA CF 555 | Biotium | 29076 | 1000 |
| Goat anti-Rabbit IgG (H+L) Cross-Adsorbed Secondary Antibody, Alexa Fluor 488 | Invitrogen | A-11008 | 500 |
| Goat anti-Chicken IgY (H+L) Secondary Antibody, Alexa Fluor 488 | Invitrogen | A-11039 | 500 |
| Goat anti-Mouse IgG (H+L) Cross-Adsorbed Secondary Antibody, Alexa Flour 647 | Invitrogen | A-21235 | 500 |
| Goat anti-Rabbit IgG (H+L) Cross-Adsorbed Secondary Antibody, Alexa Fluor 647 | Invitrogen | A-21244 | 500 |

| | | | |
|---|---|---|---|
| Donkey anti-Sheep IgG (H+L) Cross-Adsorbed Secondary Antibody, Alexa Fluor 647 | Invitrogen | A-21448 | 500 |
| NucSpot Live 650 Nuclear Stain | Biotium | 40082-T | 250 |
| Cellbrite steady 550 membrane | Biotium | 30107-T | 1000 |

**Supplementary Table S2.** Data for Figure 4G and H. Manual counting of transduced cells for different serotypes. Three image layers of z-stacks for different AAV serotypes were manually annotated by human annotators (layer 100 (25 µm), layer 200 (50 µm) and layer 300 (75 µm). The total number of cells and the number of not transduced cells was counted by examining the membrane dye channel (CellBrite550). The number of transduced cells were then calculated by subtracting the non-transduced from the transduced cells. This data can was then averaged over the layers in Figure 4G and H.

| Serotype | Layer | #Cells | #Transduced | #Not transduced | Transduction efficiency [%] |
|---|---|---|---|---|---|
| **Human annotator 1** | | | | | |
| AAV2.NN | 100 | 1254 | 989 | 265 | 79 |
| | 200 | 1391 | 1130 | 261 | 81 |
| | 300 | 648 | 451 | 197 | 70 |
| AAV9.NN | 100 | 1776 | 364 | 1412 | 20 |
| | 200 | 1666 | 406 | 1260 | 24 |
| | 300 | 797 | 226 | 571 | 28 |
| AAV2.7m8 | 100 | 915 | 116 | 799 | 13 |
| | 200 | 919 | 99 | 820 | 11 |
| | 300 | 627 | 124 | 503 | 20 |
| **Human annotator 2** | | | | | |
| AAV2.NN | 100 | 801 | 631 | 170 | 79 |
| | 200 | 445 | 375 | 70 | 84 |
| | 300 | 125 | 69 | 56 | 55 |
| **Human annotator 3** | | | | | |
| AAV2.NN | 100 | 1288 | 754 | 534 | 59 |
| | 200 | 926 | 633 | 293 | 68 |
| | 300 | 362 | 290 | 72 | 80 |

**Supplementary Table S3.** Data for Figure 4G. Automated transduction efficiency as a comparison for manual counting in Supplementary Figure 2. Three different thresholds were assessed to determine which one is best to be used in the automatized analysis pipeline. One image set of one organoid transduced each with a different serotype was manually annotated and the calculated transduction efficiency was averaged over the layers. The result was compared to the automated results in the same image stacks. The table shows the transduction efficiency resulting from the pipeline at different thresholds and the averaged transduction efficiency calculated from the manual counting averaged over the layers and its standard deviation.

| | Transduction efficiency [%] | | | | |
|---|---|---|---|---|---|
| **Serotype** | **Threshold 30 %** | **Threshold 40 %** | **Threshold 50 %** | **Human annotator** | **Standard deviation human annotator** |
| AAV2.NN | 89 | 84 | 78 | 77 | 6 |
| AAV9.NN | 34 | 24 | 16 | 24 | 4 |
| AAV2.7m8 | 20 | 15 | 12 | 14 | 5 |

**Supplementary Table S4.** Data for Figure 5C. Serotype transduction efficiency for individual organoids at different timepoints after transduction. The organoid symbols refer to the symbols used for organoid identification in Figure 5C. DIV (day in vitro) indicates the age of the organoid at the timepoint it was image. The transduction efficiency was averaged if the number of areas imaged in the same organoid was bigger than one.

| Serotype | Organoid symbol | Timepoint | DIV | Transduction efficiency (%) | Number of areas |
|---|---|---|---|---|---|
| AAV2.7m8 | circle | Td+13 | DIV244 | 38 | 1 |
| AAV2.7m8 | circle | Td+30 | DIV257 | 62 | 1 |
| AAV2.7m8 | circle | Td+40 | DIV267 | 35 | 1 |
| AAV2.7m8 | square | Td+7 | DIV235 | 31 | 1 |
| AAV2.7m8 | square | Td+30 | DIV258 | 83 | 1 |
| AAV2.7m8 | square | Td+40 | DIV268 | 66 | 1 |
| AAV2.7m8 | triangle pointing left | Td+7 | DIV236 | 5 | 1 |
| AAV2.7m8 | star | Td+7 | DIV237 | 2 | 1 |
| AAV2.7m8 | star | Td+30 | DIV260 | 64 | 2 |
| AAV2.7m8 | star | Td+40 | DIV270 | 27 | 1 |
| AAV2.7m8 | cross | Td+7 | DIV238 | 14 | 2 |
| AAV2.7m8 | cross | Td+30 | DIV261 | 20 | 1 |
| AAV2.7m8 | cross | Td+40 | DIV271 | 37 | 2 |
| AAV2.7m8 | pentagon | Td+7 | DIV239 | 26 | 1 |
| AAV2.7m8 | pentagon | Td+13 | DIV245 | 42 | 1 |
| AAV2.7m8 | pentagon | Td+30 | DIV262 | 56 | 2 |
| AAV2.7m8 | pentagon | Td+40 | DIV272 | 44 | 2 |
| AAV2.7m8 | triangle pointing right | Td+7 | DIV240 | 14 | 3 |
| AAV2.7m8 | triangle pointing right | Td+13 | DIV246 | 36 | 2 |
| AAV2.7m8 | triangle pointing right | Td+30 | DIV263 | 41 | 1 |
| AAV2.7m8 | diamond | Td+7 | DIV241 | 8 | 2 |
| AAV2.7m8 | diamond | Td+13 | DIV247 | 28 | 2 |
| AAV2.7m8 | diamond | Td+30 | DIV264 | 38 | 2 |
| AAV2.7m8 | diamond | Td+40 | DIV274 | 38 | 2 |
| AAV2.NN | circle | Td+7 | DIV234 | 99 | 1 |
| AAV2.NN | circle | Td+13 | DIV244 | 78 | 2 |
| AAV2.NN | circle | Td+30 | DIV257 | 98 | 1 |
| AAV2.NN | circle | Td+40 | DIV267 | 90 | 1 |
| AAV2.NN | square | Td+7 | DIV235 | 82 | 1 |
| AAV2.NN | square | Td+30 | DIV258 | 98 | 1 |
| AAV2.NN | square | Td+40 | DIV268 | 95 | 1 |
| AAV2.NN | triangle pointing left | Td+7 | DIV236 | 69 | 2 |
| AAV2.NN | triangle pointing left | Td+30 | DIV259 | 96 | 2 |
| AAV2.NN | triangle pointing left | Td+40 | DIV269 | 62 | 2 |
| AAV2.NN | star | Td+7 | DIV237 | 90 | 1 |
| AAV2.NN | star | Td+30 | DIV260 | 79 | 2 |
| AAV2.NN | cross | Td+7 | DIV238 | 90 | 1 |
| AAV2.NN | cross | Td+30 | DIV261 | 51 | 2 |
| AAV2.NN | cross | Td+40 | DIV271 | 38 | 1 |
| AAV2.NN | pentagon | Td+7 | DIV239 | 84 | 1 |
| AAV2.NN | pentagon | Td+13 | DIV245 | 88 | 1 |
| AAV2.NN | pentagon | Td+30 | DIV262 | 99 | 1 |
| AAV2.NN | pentagon | Td+40 | DIV272 | 98 | 1 |
| AAV2.NN | diamond | Td+7 | DIV241 | 57 | 2 |
| AAV2.NN | diamond | Td+13 | DIV247 | 79 | 3 |
| AAV2.NN | diamond | Td+30 | DIV264 | 86 | 1 |
| AAV9.NN | circle | Td+7 | DIV234 | 24 | 1 |

| | | | | | |
|---|---|---|---|---|---|
| AAV9.NN | circle | Td+13 | DIV244 | 40 | 2 |
| AAV9.NN | circle | Td+30 | DIV257 | 26 | 1 |
| AAV9.NN | circle | Td+40 | DIV267 | 84 | 2 |
| AAV9.NN | square | Td+7 | DIV235 | 35 | 1 |
| AAV9.NN | square | Td+30 | DIV258 | 66 | 1 |
| AAV9.NN | square | Td+40 | DIV268 | 79 | 1 |
| AAV9.NN | triangle pointing left | Td+7 | DIV236 | 15 | 2 |
| AAV9.NN | triangle pointing left | Td+30 | DIV259 | 55 | 1 |
| AAV9.NN | triangle pointing left | Td+40 | DIV269 | 26 | 1 |
| AAV9.NN | star | Td+7 | DIV237 | 7 | 1 |
| AAV9.NN | star | Td+40 | DIV270 | 30 | 1 |
| AAV9.NN | cross | Td+7 | DIV238 | 11 | 2 |
| AAV9.NN | cross | Td+30 | DIV261 | 27 | 1 |
| AAV9.NN | cross | Td+40 | DIV271 | 83 | 1 |
| AAV9.NN | pentagon | Td+7 | DIV239 | 33 | 1 |
| AAV9.NN | pentagon | Td+13 | DIV245 | 35 | 2 |
| AAV9.NN | pentagon | Td+30 | DIV262 | 58 | 1 |
| AAV9.NN | pentagon | Td+40 | DIV272 | 70 | 1 |
| AAV9.NN | triangle pointing right | Td+7 | DIV240 | 11 | 2 |
| AAV9.NN | triangle pointing right | Td+13 | DIV246 | 37 | 2 |
| AAV9.NN | diamond | Td+13 | DIV247 | 62 | 1 |
| AAV9.NN | diamond | Td+30 | DIV264 | 49 | 1 |

**Supplementary Table S5.** Data for figure 5C. Mean serotype transduction efficiency at different timepoints after transduction.

| Serotype | Timepoint | Mean transduction efficiency [%] | Standard error of the mean [%] |
|---|---|---|---|
| AAV2.7m8 | Td+7 | 14 | 5 |
| AAV2.7m8 | Td+13 | 36 | 3 |
| AAV2.7m8 | Td+30 | 52 | 8 |
| AAV2.7m8 | Td+40 | 41 | 6 |
| AAV9.NN | Td+7 | 19 | 5 |
| AAV9.NN | Td+13 | 43 | 7 |
| AAV9.NN | Td+30 | 47 | 7 |
| AAV9.NN | Td+40 | 62 | 12 |
| AAV2NN | Td+7 | 82 | 6 |
| AAV2NN | Td+13 | 82 | 4 |
| AAV2NN | Td+30 | 87 | 7 |
| AAV2NN | Td+40 | 77 | 12 |

**Supplementary Table S6.** Data for figure 6A. Mean serotype transduction efficiency at different depths on Td+7.

| Serotype | Bin [µm] | Mean transduction efficiency [%] | Standard error of the mean [%] |
|---|---|---|---|
| AAV2.7m8 | 0-10 | 15 | 10 |
| AAV2.7m8 | 10-20 | 9 | 6 |
| AAV2.7m8 | 20-30 | 11 | 4 |
| AAV2.7m8 | 30-40 | 11 | 4 |
| AAV2.7m8 | 40-50 | 13 | 4 |
| AAV2.7m8 | 50-60 | 19 | 7 |
| AAV2.7m8 | 60-70 | 20 | 7 |
| AAV2.7m8 | 70-80 | 22 | 8 |

| | | | |
|---|---|---|---|
| AAV2.7m8 | 80-90 | 20 | 8 |
| AAV2.7m8 | 90-100 | 16 | 7 |
| AAV9.NN | 0-10 | 12 | 7 |
| AAV9.NN | 10-20 | 18 | 5 |
| AAV9.NN | 20-30 | 19 | 5 |
| AAV9.NN | 30-40 | 22 | 6 |
| AAV9.NN | 40-50 | 22 | 6 |
| AAV9.NN | 50-60 | 24 | 6 |
| AAV9.NN | 60-70 | 23 | 6 |
| AAV9.NN | 70-80 | 20 | 5 |
| AAV9.NN | 80-90 | 17 | 6 |
| AAV9.NN | 90-100 | 11 | 5 |
| AAV2NN | 0-10 | 50 | 18 |
| AAV2NN | 10-20 | 56 | 14 |
| AAV2NN | 20-30 | 71 | 8 |
| AAV2NN | 30-40 | 77 | 6 |
| AAV2NN | 40-50 | 82 | 7 |
| AAV2NN | 50-60 | 90 | 5 |
| AAV2NN | 60-70 | 91.9 | 2.9 |
| AAV2NN | 70-80 | 93.1 | 2.1 |
| AAV2NN | 80-90 | 93.8 | 2.2 |
| AAV2NN | 90-100 | 87 | 8 |

**Supplementary Movie S1:** Confocal image and segmentation of a 226 days old human retinal organoid transduced with AAV2.7m8-egfp**.** The cytoplasmic eGFP is displayed in cyan, the stained nuclei in magenta and the nuclei segmentation in grey-scale. The ellipsoidal approximation of nuclei belonging to transduced cells is displayed in a color encoding the z position (spectral gradient) in the retina.
